# Chromosomal-level genome assembly of the long-spined sea urchin *Diadema setosum* (Leske, 1778)

**DOI:** 10.1101/2024.01.16.575490

**Authors:** Hong Kong Biodiversity Genomics Consortium, Project Coordinator and Co-Principal Investigators, Jerome H.L. Hui, Ting Fung Chan, Leo L. Chan, Siu Gin Cheung, Chi Chiu Cheang, James K.H. Fang, Juan D. Gaitan-Espitia, Stanley C.K. Lau, Yik Hei Sung, Chris K.C. Wong, Kevin Y.L. Yip, Yingying Wei, DNA extraction, library preparation and sequencing, Wai Lok So, Genome assembly and gene model prediction, Wenyan Nong, Sample collector, animal culture and logistics, Apple P.Y. Chui, Thomas H.W. Fong, Ho Yin Yip

**Affiliations:** School of Life Sciences, Simon F.S. Li Marine Science Laboratory, State Key Laboratory of Agrobiotechnology, Institute of Environment, Energy and Sustainability, The Chinese University of Hong Kong, Hong Kong, China; School of Life Sciences, State Key Laboratory of Agrobiotechnology, The Chinese University of Hong Kong, Hong Kong SAR, China; State Key Laboratory of Marine Pollution and Department of Biomedical Sciences, City University of Hong Kong, Hong Kong SAR, China; State Key Laboratory of Marine Pollution and Department of Chemistry, City University of Hong Kong, Hong Kong SAR, China; Department of Science and Environmental Studies, The Education University of Hong Kong, Hong Kong SAR, China; EcoEdu PEI, Charlottetown, PE, C1A 4B7, Canada; Department of Food Science and Nutrition, Research Institute for Future Food, and State Key Laboratory of Marine Pollution, The Hong Kong Polytechnic University, Hong Kong SAR, China; The Swire Institute of Marine Science and School of Biological Sciences, The University of Hong Kong, Hong Kong SAR, China; Department of Ocean Science, The Hong Kong University of Science and Technology, Hong Kong SAR, China; Science Unit, Lingnan University, Hong Kong SAR, China; School of Allied Health Sciences, University of Suffolk, Ipswich, IP4 1QJ, UK; Croucher Institute for Environmental Sciences, and Department of Biology, Hong Kong Baptist University, Hong Kong SAR, China; Department of Computer Science and Engineering, The Chinese University of Hong Kong, Hong Kong SAR, China; Sanford Burnham Prebys Medical Discovery Institute, La Jolla, CA, USA; Department of Statistics, The Chinese University of Hong Kong, Hong Kong SAR, China; School of Life Sciences, The Chinese University of Hong Kong, Hong Kong SAR, China

## Abstract

The long-spined sea urchin *Diadema setosum* is an algal and coral feeder widely distributed in the Indo-Pacific and can cause severe bioerosion on the reef community. Nevertheless, the lack of genomic information has hindered the study its ecology and evolution. Here, we report the chromosomal-level genome (885.8 Mb) of the long-spined sea urchin *D. setosum* using a combination of PacBio long-read sequencing and Omni-C scaffolding technology. The assembled genome contained scaffold N50 length of 38.3 Mb, 98.1 % of BUSCO (Geno, metazoa_odb10) genes, and with 98.6% of the sequences anchored to 22 pseudo-molecules/chromosomes. A total of 27,478 genes including 23,030 protein-coding genes were annotated. The high-quality genome of *D. setosum* presented here provides a significant resource for further understanding on the ecological and evolutionary studies of this coral reef associated sea urchin.

## Introduction

Similar to other echinoderms, sea urchins lack the vertebral column and can metamorphose from juvenile bilateral swimming larvae into radial symmetrical adults (Li et al., 2020; Lowe et al., 2015). Owing to its critical phylogenetic position, sea urchin offers the understanding of how deuterostomes evolved (e.g. Odekunle et al., 2019; Yañez-Guerra et al., 2020; Sakai et al., 2020; Logeswaran et al. 2021; Zheng et al., 2022). To date, 16 sea urchin genomes are available according to the data presented on NCBI (25 October 2023) and only five of them (all in the order Camarodonta) are assembled in chromosomal level (*Lytechinus variegatus* (Davidson et al., 2020), *Lytechinus pictus* (Warner et al., 2021), *Heliocidaris erythrogramma* (Davidson et al., 2022), *Heliocidaris tuberculata* (Davidson et al., 2022) and *Echinometra lucunter* (Davidson et al., 2023).

*Diadema setosum* (Leske, 1778) in the order Diadematoida, commonly known as the porcupine or long-spined sea urchin, is considered as one of the oldest known extant species in the genus *Diadema* (Coppard & Campbell, 2006). *D. setosum* displays features of typical sea urchin, including a dorso-ventrally compressed body and equipped with particularly long, brittle and hollow spines that are mildly venomous (Bilecenoğlu et al., 2019; Voulgaris et al., 2021). This species can be easily differentiated from other *Diadema* species by the presence of five distinctive white dots at the aboral side around the anal pore between the ambulacral grooves (Figure 1A). Sexually matured individuals have been documented to have an average weight from 35 to 80 g and an average test size ranges from 7 to 8 cm in diameter and approximately 4 cm in height (Alsaffar & Lone, 2000; Coppard & Campbell, 2006). Due to its high invasiveness to localities beyond its natural range, *D. setosum* is now widely distributed in the tropical regions throughout the Indo-Pacific basin and can now be found latitudinally from Japan to Africa and longitudinally from the Red Sea to Australia (Öndes et al., 2022).

**Figure 1.**
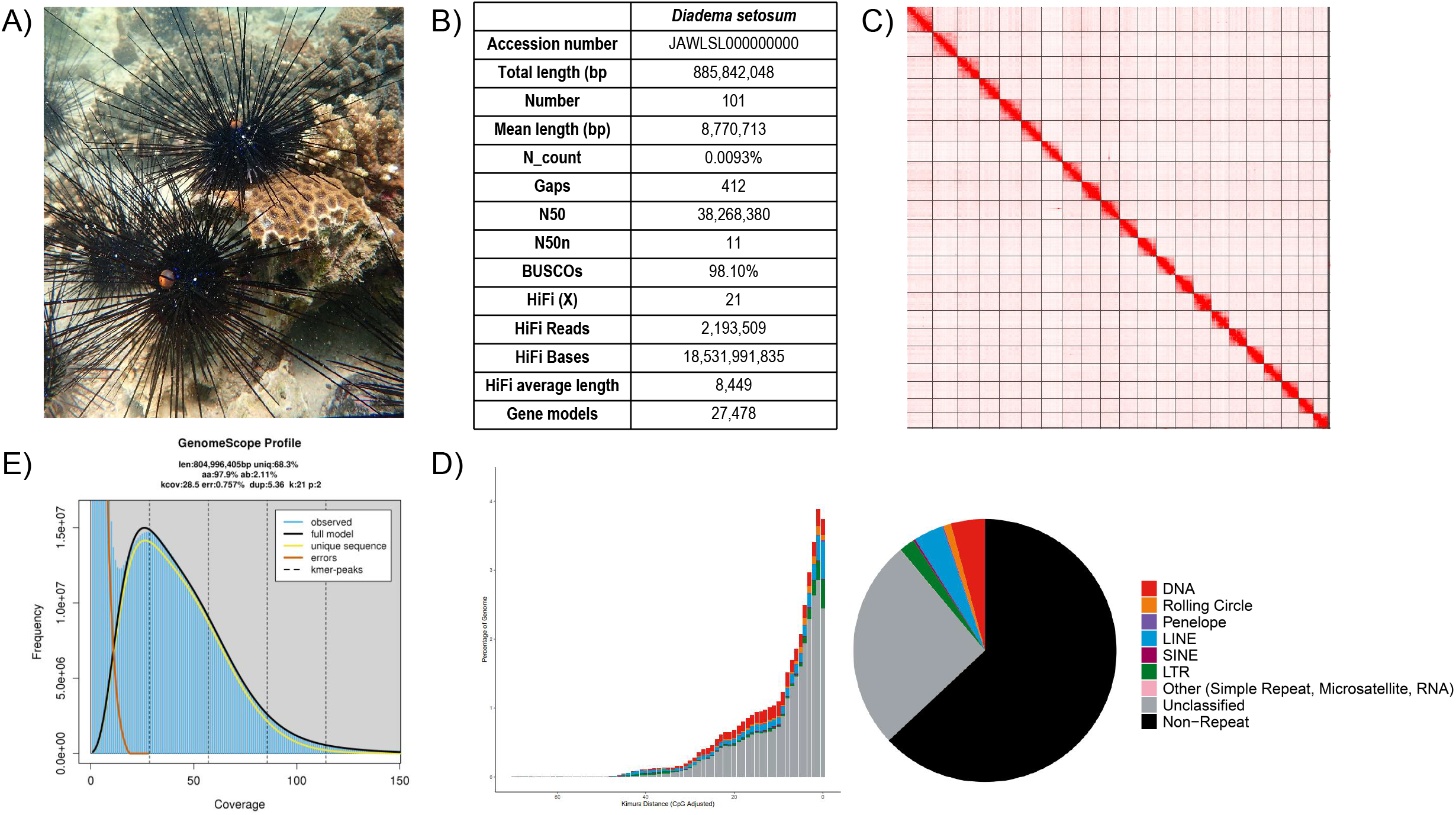
Genomic information of *Diadema setosum*. A) Photo of *D. setosum*; B) Statistics of the assembled genome; C) Omni-C contact map of the assembly visualised using Juicebox v1.11.08 (details can be found in Additional Table 6); D) Genomescope report with k-mer = 21; E) Repetitive elements distribution in the assembled genome.

*D. setosum* can live up as deep as 70 m below the sea level and is usually reef-associated (Lane et al., 2001). It is a prolific grazer that feeds on the macroalgae that can be found on the surface of various substrata, as well as the algae that are associated with the coral skeleton (Bak, 1994; Carreiro-Silva and McClanahan, 2001). Whilst a normal level of grazing eliminates competitive algae and can potentially offer a more suitable environments for coral settlement and development, overgrazing results in bioerosion, which thus in turn deteriorates the reef ecosystem and a reduction in the complexity of coral community (Carpenter and Edmunds, 2006; Dumont et al., 2013). Furthermore, overpopulated sea urchins can reduce coral recruitment and the growth of juvenile coral can be hindered (Bak and van Eys, 1975; Sammarco, 1980; Davies and Vize, 2008).

## Context

Here, we report a high-quality genome assembly of *D. setosum* in the order Diadematoida and family Diadematidae.

## Methods

### Collection and husbandry of samples

The long-spined sea urchin, *Diadema setosum*, was collected at the coastal area of the Tolo Habour in Hong Kong in November 2022. The animals were maintained at 35 ppt artificial sea water alive at 23°C until DNA and RNA isolation. Animals were fed with frozen clams or shrimps once a week.

### Isolation of high molecular weight genomic DNA, quantification, and qualification

High molecular weight genomic DNA was isolated from a single individual. The urchin was first removed from the culture and the test was opened with a pair of scissors. The internal tissue, except the gut, was snap-frozen in liquid nitrogen and ground to fine powder. DNA extraction was performed with the Qiagen MagAttract HMW kit (Qiagen Cat. No. 67563) following the manufacturer’s protocol. In brief, 1 g of powdered sample was put in a microcentrifuge tube together with 200 μl 1X PBS and RNase A, Proteinase K, and Buffer AL were added to the tube subsequently. The mixture was incubated at room temperature (∼22 °C) for 3 hours. The sample was then eventually eluted with 120 μl of elution buffer (PacBio Ref. No. 101-633-500). Throughout the extraction progress, wide-bore tips were used whenever DNA was transferred. The eluted sample was quantified by the Qubit^®^ Fluorometer and Qubit^™^ dsDNA HS and BR Assay Kits (Invitrogen^™^ Cat. No. Q32851). In total, 10 μg of DNA was collected for subsequent steps. The purity of the sample was examined by the NanoDrop^™^ One/OneC Microvolume UV-Vis Spectrophotometer, with the standard A260/A280: ∼1.8 and A260/A230: >2.0. The quality and the fragment distribution of the isolated genomic DNA was examined by the overnight pulse-field gel electrophoresis, together with three DNA markers (λ-Hind III digest, Takara Cat. No. 3403; DL15,000 DNA Marker, Takara Cat. No. 3582A and CHEF DNA Size Standard-8-48 kb Ladder, Cat. No. 170-3707). The DNA was then diluted in elution buffer to prepare a 300 ng solution for gel electrophoresis and the electrophoresis profile was set as follow: 5k as the lower end and 100k as the higher end for the molecular weight; Gradient=6.0V/cm; Run time = 15 h:16 min; Included angle = 120 °; Int. Sw. Tm = 22 s; Fin. Sw. Tm = 0.53 s; Ramping factor: a = Linear. The gel was run in 1.0% PFC agarose in 0.5X TBE buffer at 14 °C.

### DNA shearing, library preparation and sequencing

10 μg of *D. setosum* DNA in 120 μl elution buffer was transferred to a g-tube (Covaris Part No. 520079). The sample was then proceeded to 6 passes of centrifugation with 2,000 xg for 2 min. The sheared DNA was collected with a 2 ml DNA LoBind^®^ Tube (Eppendorf Cat. No. 022431048) at 4 °C until library preparation. Overnight pulse-field gel electrophoresis was performed to examine the fragment distribution of the sheared DNA, with the same electrophoresis profile as described in the last section.

A SMRTbell library was then constructed with the SMRTbell® prep kit 3.0 (PacBio Ref. No. 102-141-700), following the manufacturer’s protocol. In brief, the sheared DNA was first subjected to DNA repair, and both ends of each DNA strand was polished and tailed with an A-overhang. Ligation of T-overhang SMRTbell adapters was then performed and the SMRTbell library was purified with SMRTbell^®^ cleanup beads (PacBio Ref. No. 102158-300). The concentration and size of the library were examined with the Qubit^®^ Fluorometer and Qubit™ dsDNA HS and BR Assay Kits (Invitrogen^™^ Cat. No. Q32851), and an overnight pulse-field gel electrophoresis, respectively. A nuclease treatment step was performed afterwards to remove any non-SMRTbell structures in the library and a final size-selection step was performed to remove the short fragments in the library with 35% AMPure PB beads.

The Sequel^®^ II binding kit 3.2 (PacBio Ref. No. 102-194-100) was used for final preparation of sequencing. In brief, Sequel II primer 3.2 and Sequel II DNA polymerase 2.2 were annealed and bound to the SMRTbell structures in the library. The library then was loaded at an on-plate concentration of 90 pM using the diffusion loading mode. The sequencing was performed on the Sequel IIe System with an internal control provided in the binding kit. The sequencing was prepared and run in 30-hour movies, with 120 min pre-extension. The movie was captured by the software SMRT Link v11.0 (PacBio) and HiFi reads are generated and collected for further analysis. In total, one SMRT cell was used in the sequencing. Details of the sequencing data are listed in Supplementary Information 1.

### Omni-C library preparation and sequencing

An Omni-C library was constructed using the Dovetail® Omni-C® Library Preparation Kit (Dovetail Cat. No. 21005) according to the manufacturer’s instructions. In brief, 60 mg of frozen powered tissue sample was added into 1 mL 1X PBS, where the genomic DNA was crosslinked with formaldehyde and the DNA was then digested with endonuclease DNase I. Subsequently, the concentration and fragment size of the digested sample was validated by the Qubit® Fluorometer and Qubit™ dsDNA HS and BR Assay Kits (Invitrogen™ Cat. No. Q32851), and the TapeStation D5000 HS ScreenTape, respectively. Afterwards, both ends of the DNA were polished and a biotinylated bridge adaptor was ligated at 22 °C for 30 min. Proximity ligation between crosslinked DNA fragments was then performed at 22 °C for 1 hour, followed by the reverse crosslinked of DNA, and it is then purified with SPRIselect™ Beads (Beckman Coulter Product No. B23317).

End repair and adapter ligation were performed with the Dovetail™ Library Module for Illumina (Dovetail Cat. No. 21004). In brief, DNA was tailed with an A-overhang, and ligate with Illumina-compatible adapters at 20 °C for 15 min. The Omni-C library was then sheared into small fragments with USER Enzyme Mix and purified with SPRIselect™ Beads. Subsequently, DNA fragments were isolated with Streptavidin Beads. Universal and Index PCR Primers from the Dovetail™ Primer Set for Illumina (Dovetail Cat. No. 25005) were used to amplify the DNA library. A final size selection step was completed with SPRIselect™ Beads with DNA fragments ranging between 350 bp and 1000 bp only. The concentration and fragment size of the sequencing library was assessed by the Qubit® Fluorometer and Qubit™ dsDNA HS and BR Assay Kits, and the TapeStation D5000 HS ScreenTape, respectively. Qualified library was sequenced on an Illumina HiSeq-PE150 platform. Details of the sequencing data are listed in Supplementary Information 1.

### RNA extraction and transcriptome sequencing

Total RNA from the internal tissues were isolated from one individual using the TRIzol reagent (Invitrogen) following the manufacturer’s protocol. The quality of the extracted RNA was validated with the NanoDrop™ One/OneC Microvolume UV-Vis Spectrophotometer (Thermo Scientific™ Cat. No. ND-ONE-W) and 1% agarose gel electrophoresis. The qualified samples were sent to Novogene Co. Ltd (Hong Kong, China) for polyA selected RNA sequencing library construction using the TruSeq RNA Sample Prep Kit v2 (Illumina Cat. No. RS-122-2001), and 150 bp paired-end (PE) sequencing. Agilent 2100 Bioanalyser (Agilent DNA 1000 Reagents) was used to measure the insert size and concentration of the final library. Details of the sequencing data are shown in Supplementary Information 1.

### Genome assembly and gene model prediction

*De novo* genome assembly was completed using Hifiasm (Cheng et al., 2021), and the Hifiasm output assembly was BLAST to the NT database and the BLAST output was used as input for BlobTools (v1.1.1) (Laetsch & Blaxter, 2017) to validate and remove any possible contaminations (Additional Information 1). Haplotypic duplications were detected and removed using purge_dups according to the depth of HiFi reads (Guan et al., 2020). Proximity ligation data from the Omni-C library were used to scaffold the PacBio genome by YaHS (Zhou et al., 2022). A kmer-based statistical method was performed to estimate the heterozygosity, while repeat content and size were analyzed by Jellyfish (Marçais & Kingsford, 2011) and GenomeScope (http://qb.cshl.edu/genomescope/genomescope2.0/) (Ranallo-Benavidez et al., 2020). Transposable elements (TEs) were annotated, using the automated Earl Grey TE annotation pipeline (version 1.2, https://github.com/TobyBaril/EarlGrey) (Baril et al., 2022). Mitochondrial genome was assembled using MitoHiFi (v2.2, https://github.com/marcelauliano/MitoHiFi) (Allio et al., 2020).

For gene model prediction, RNA sequencing data was first processed with Trimmomatic (Bolger, Lohse & Usadel, 2014) and transformed to transcripts using genome-guided Trinity (Grabherr et al., 2011). AUGUSTUS (Stanke et al., 2006) was trained using BUSCO (Manni et al., 2021), while GeneMark-ES (Ter-Hovhannisyan et al 2008) was used for *ab initio* gene prediction. Gene models were then predicted by funannotate using the parameters “--repeats2evm --protein_evidence uniprot_sprot.fasta --genemark_mode ET -- optimize_augustus --organism other --max_intronlen 350000”. The gene models from several prediction sources including GeneMark, high-quality Augustus predictions (HiQ), pasa, Augustus, GlimmerHM and snap were passed to Evidence Modeler to generate the annotation files. PASA was employed to update the EVM consensus predictions. In addition, UTR annotations were added and models for alternatively spliced isoforms were created.

## Results

A total of 18.5 Gb of HiFi bases were generated with an average HiFi read length of 8,449 bp with 21x data coverage (Supplementary Information 1). The assembled genome size was 885.8 Mb, with 101 scaffolds and a scaffold N50 of 38.3 Mb in 11 scaffolds and the complete BUSCO estimation to be 98.1% (metazoa_odb10) (Figure 1B; Table 1). By incorporating 67.5 Gb Omni-C data, the assembly anchored 98.6% of the scaffolds into 22 pseudochromosomes, which matches the karyotype of *D. setosum* (2n = 44) (Shingaki & Uehara, 1984) (Figure 1C; Table 2). The assembled *D. setosum* genome size is comparable to other published sea urchin genomes (Davidson et al., 2020; Warner et al., 2021; Davidson et al., 2022; Davidson et al., 2023) and to the estimated size of 804 Mb by GenomeScope with a 2.11% heterozygosity rate (Figure 1D; Supplementary Information 2). Moreover, telomeric repeats were identified in 16 out of 22 pseudochromosomes (Table 3).

Total RNA sequencing data was also obtained from a single *D. setosum* individual. The final assembled transcriptome contained 135,063 transcripts, with 113,391 Trinity annotated genes (with an average length of 838 bp and a N50 length of 1,456 bp), and was used to perform gene model prediction. A total of 27,478 gene models were generated with 23,030 predicted protein-coding genes, with a mean of 483 bp coding sequence length (Figure 1B; Table 1).

For repeat elements, a total repetitive content of 36.98% was identified in the assembled genome including 25.87% unclassified elements (Figure 1E; Table 4). Among the known repeats, DNA is the most abundant (4.18%), followed by LINE elements (3.64%) and LTR (1.92%), whereas Rolling Circle, SINE, Penelope and other are only present in low proportions (Rolling Circle: 0.92%, SINE: 0.23%, Penelope: 0.17%, other: 0.04%).

### Conclusion and future perspectives

In sum, the high quality chromosomal-level genome assembly of *D. setosum* presented here provides further understanding of the ecology and evolution of sea urchins.

## Data Records

The final genome assembly were deposited to NCBI under accession numbers GCA_033980235.1 (https://www.ncbi.nlm.nih.gov/datasets/genome/GCA_033980235.1/). The raw reads generated in this study were submitted to the NCBI database under the BioProject accessions PRJNA973839 (https://www.ncbi.nlm.nih.gov/bioproject/PRJNA973839). The genome and genome annotation files, as well as an Additional Table and an Additional Information were also deposited in figshare (https://doi.org/10.6084/m9.figshare.24408319).

## Data validation and quality control

Quality check of samples in DNA extraction and PacBio library preparation were processed with NanoDrop^™^ One/OneC Microvolume UV-Vis Spectrophotometer, Qubit^®^ Fluorometer, and overnight pulse-field gel electrophoresis. The Omni-C library was subjected to quality check by Qubit® Fluorometer and TapeStation D5000 HS ScreenTape.

For genome assembly, the validation of contamination scaffolds from the Hifiasm output was done by searching at the NT database through BLAST, from which the resultant output was analysed by BlobTools (v1.1.1) (Laetsch & Blaxter, 2017) (Supplementary Information 3). Furthermore, a kmer-based statistical approach was performed to estimate the genome heterozygosity. The repeat content and their size were estimated by Jellyfish (Marçais & Kingsford, 2011) and GenomeScope (Figure 1E and Additional Table 5) (http://qb.cshl.edu/genomescope/genomescope2.0/) (Ranallo-Benavidez et al., 2020). Benchmarking Universal Single-Copy Orthologs (BUSCO, v5.5.0) (Manni et al 2021) was run to evaluate the completeness of the genome assembly and gene annotation with the metazoan dataset (metazoa_odb10).

## Author’ contribution

JHLH, TFC, LLC, SGC, CCC, JKHF, JDG, SCKL, YHS, CKCW, KYLY and YW conceived and supervised the study; APYC and THWF collected the sea urchin samples; HYY maintained the animal culture; WLS performed DNA extraction, library preparation and genome sequencing; WN carried out genome assembly and gene model prediction.

---

**Table 1**. Genome assembly statistic and sequencing information.

**Table 2**. Scaffold information of 22 pseudochromosomes.

**Table 3**. List of telomeric repeats identified in the genome.

**Table 4**. Statistics of annotated repetitive elements.

**Supplementary Information 1**. Genome and transcriptome sequencing information.

**Supplementary Information 2**. GenomeScope statistics with K-mer length 21.

**Supplementary Information 3**. Genome assembly QC and contaminant/cobiont detection.

