## Supplementary material for "Chromosomal-level genome assembly of the long-spined sea urchin *Diadema setosum* (Leske, 1778)": EBPHK_Sea urchin Supplementary Information.docx

**Supplementary Information 1.** Genome and transcriptome sequencing information.

| **Genome sequencing data** | | | | |
| --- | --- | --- | --- | --- |
| **Liabrary** | **No. of reads** | **No. of bases** | **Coverage(X)** | **Accesion** |
| PacBio HiFi | 2,193,509 | 18,531,991,835 | 21 | SRR24631719 |
| Omnic | 450,451,192 | 67,567,678,800 | 76 | SRR26502301 |
| **Transcriptome sequencing data** | | | | |
| **Sample name** | **No. of reads** | **No. of bases** | **Accesion** | |
| DseRNA | 40,875,262 | 6,131,231,889 | SRR24694066 | |

**Supplementary Information 2.** GenomeScope statistics with K-mer length 21.

| Property | min | max |
| --- | --- | --- |
| Homozygous (aa) | 97.85% | 97.93% |
| Heterozygous (ab) | 2.07% | 2.15% |
| Genome Haploid Length (bp) | 795,402,175 | 804,996,405 |
| Genome Repeat Length (bp) | 251,904,808 | 254,943,312 |
| Genome Unique Length (bp) | 543,497,367 | 550,053,093 |
| Model Fit | 75.49% | 98.83% |
| Read Error Rate | 0.76% | 0.76% |

**Supplementary Information 3.** Genome assembly QC and contaminant/cobiont detection.

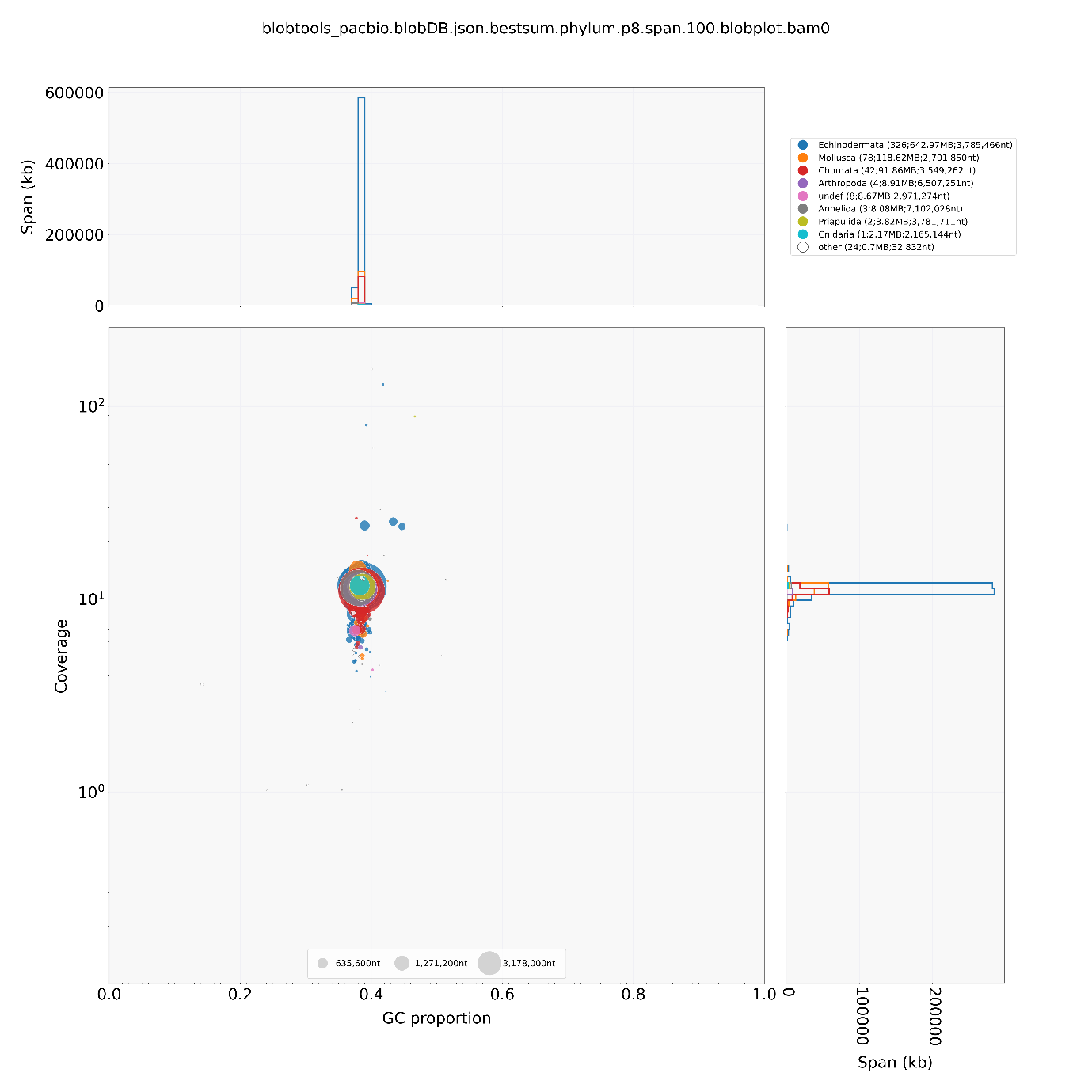

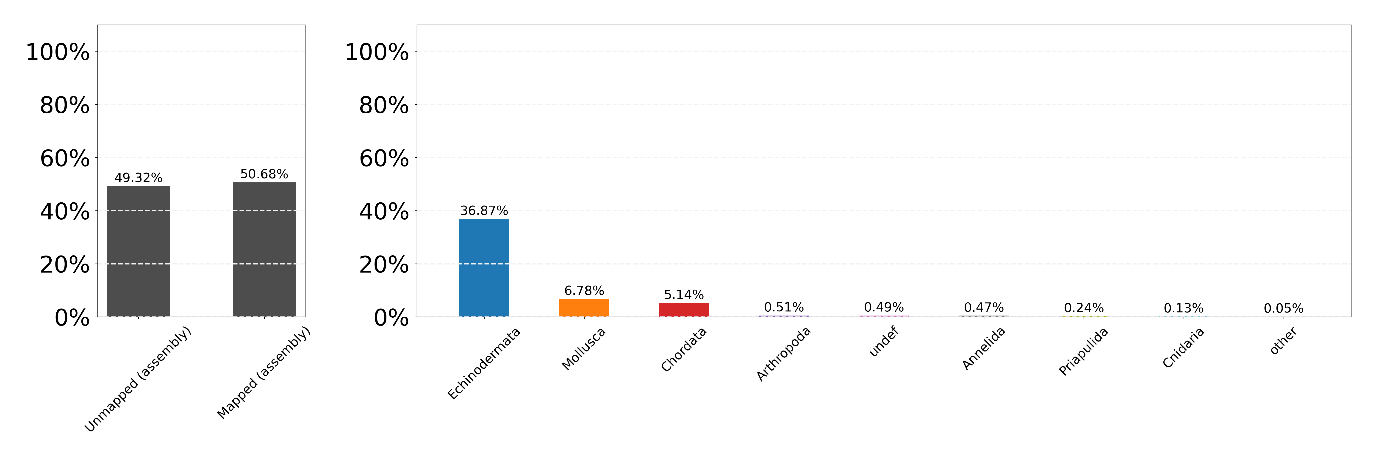
